# *Plasmodium falciparum* protein phosphatase PP7 is required for early ring-stage development

**DOI:** 10.1101/2024.04.08.588616

**Authors:** Avnish Patel, Aline Fréville, Joshua A. Rey, Helen R. Flynn, Konstantinos Koussis, Mark J. Skehel, Michael J. Blackman, David A. Baker

## Abstract

We previously reported that the *Plasmodium falciparum* putative serine/threonine protein phosphatase 7 (PP7) is a high confidence substrate of the cAMP-dependent protein kinase (PKA). Here we explore the function of PP7 in asexual *P. falciparum* blood stage parasites. We show that conditional disruption of PP7 leads to a severe growth arrest. We show that PP7 is a calcium-dependent phosphatase which interacts with calmodulin and calcium-dependent protein kinase 1 (CDPK1), consistent with a role in calcium signalling. Notably, PP7 was found to be dispensable for erythrocyte invasion, but was crucial for ring-stage development, with PP7-null parasites arresting shortly following invasion and showing no transition to ameboid forms. Phosphoproteomic analysis revealed that PP7 may regulate certain PKAc substrates. Its interaction with calmodulin and CDPK1 further emphasise a role in calcium signalling, while its impact on early ring development and PKAc substrate phosphorylation underscores its importance in parasite development.

## Introduction

Protein phosphorylation is an essential post-translational modification event that is dynamically controlled by the action of protein kinases and protein phosphatases in all eukaryotic cells. The major human malaria pathogen *Plasmodium falciparum* possesses 90 putative protein kinases (1, 2) and 27 putative protein phosphatases (3). Genome-wide screens in *P. berghei* , a rodent malaria species, has identified the stage-specific essentiality of kinases and phosphatases (4, 5). With the widespread adoption of conditional gene disruption technologies in *Plasmodium*, several of the essential kinases have been further studied for their cell-specific roles (6–11). By contrast, essential protein phosphatases have remained relatively under-explored (12), yet an improved understanding of their function is a requisite to discerning the global role of protein phosphorylation dynamics in the complex malaria parasite life cycle, and can also inform target-based drug discovery efforts for rational drug design. Previous studies have assigned functions for essential phosphatases to processes late in the asexual blood stage cycle, revealing roles for calcineurin for merozoite host cell attachment (13, 14), PP1 in schizont DNA replication and egress (15), and NIF4 in merozoite invasion (16). However, to our knowledge no malarial protein phosphatase has been shown to play a role in the early developmental phases of asexual blood stage parasites.

We previously identified 39 potential substrates of the cAMP dependent protein kinase catalytic subunit (PKAc), an essential kinase required for merozoite invasion of host erythrocytes. Phosphosites within these protein substrates were found to be upregulated following disruption of adenylyl cyclase β or PKAc, and correspondingly downregulated following deletion of phosphodiesterase β (17) (7). Amongst the putative PKAc substrates identified, PP7 was considered of particular interest in our efforts to elucidate the cell signalling events underlying erythrocyte invasion due to its predicted essentiality and its peak transcriptional expression in mature schizonts.

Here we investigate the function of *P. falciparum* PP7. We find it to be indispensable for asexual blood stage parasite growth, and specifically for the development of early ring-stage parasites. We show that PP7 is expressed in late schizont stages, that its activity is Ca^2+^-dependent, and that it interacts with both calmodulin and the Ca^2+^-dependent protein kinase CDPK1. Finally, we use global phospho-proteomics to identify sites that are differentially regulated by PP7.

## Results

### The architecture of PP7 includes calcium-binding domains

PP7 is annotated as a member of the Phospho-Protein Phosphatases (PPP) family (PlasmoDB (18) PF3D7_1423300). Its overall structure comprises a central predicted serine/threonine protein phosphatase domain (19) flanked by putative Ca^2+^-binding-related domains: an N-terminal calmodulin binding IQ domain (20) and two C-terminal Ca^2+^-binding EF hands (21) (Fig 1A). This architecture is characteristic of members of one of the five sub-families of the phosphoprotein phosphatase (PPP) family, PPEF/PP7, with EF referring to the presence of an EF-hand domain downstream of the phosphatase catalytic domain. Figure S1 shows the domain structure of the *P. falciparum* PP7 and alignments with orthologues from other eukaryotes (22).

**Figure 1.**
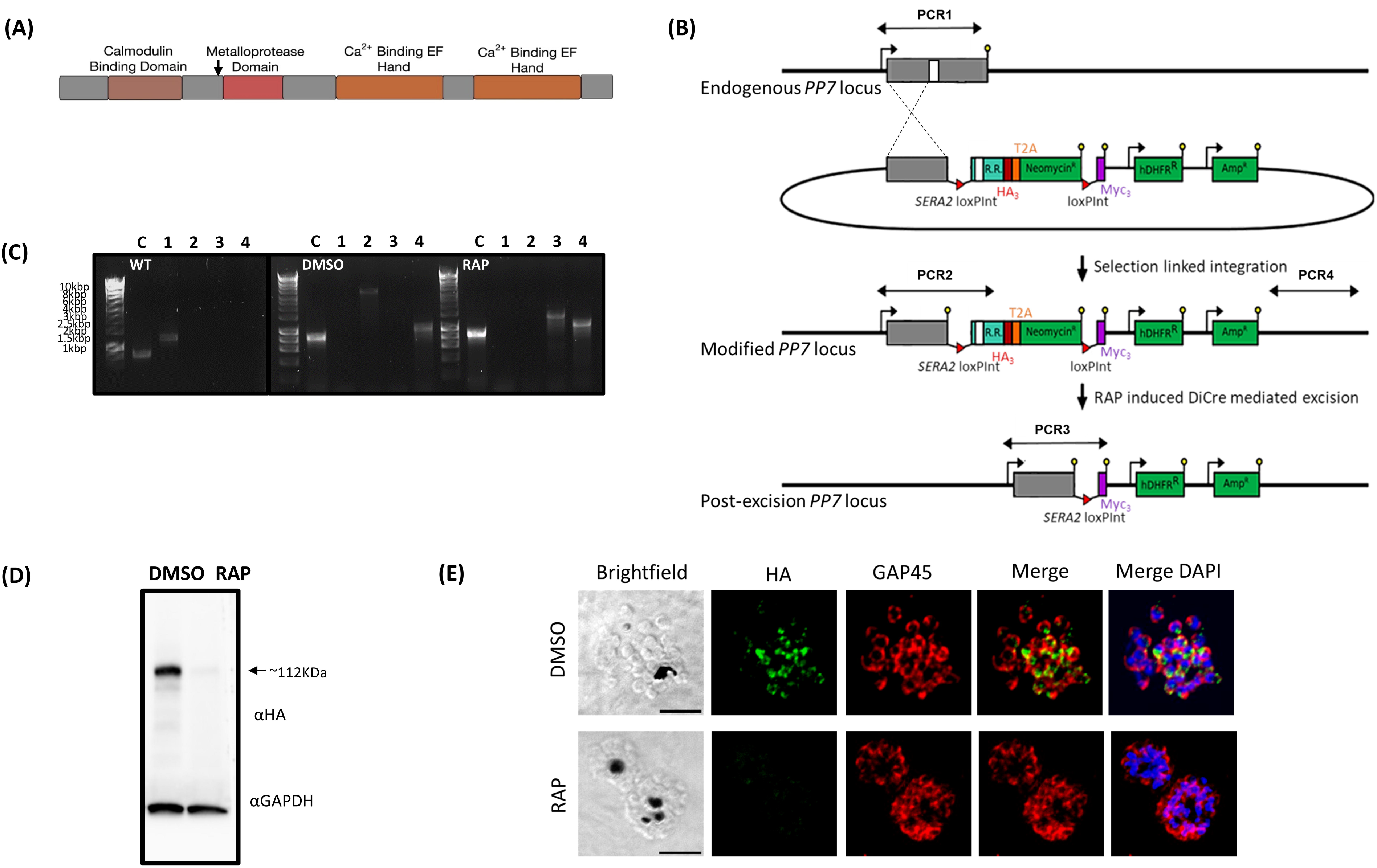
(1A) Cartoon of the predicted *P. falciparum* PP7 protein showing the positions of the individual domains. Amino acid numbers are shown in black. The arrow indicates the point at which the protein product is truncated when the modified locus is excised in the transgenic parasites. (1B) Schematic representation of the SLI strategy (39) used to produce the PP7-HA:loxP DiCre line and resultant RAP-induced disruption of the modified gene. Double-headed arrows represent the regions amplified by PCR in (1C). Red arrowheads, *loxP* sites, yellow lollipops, translational stop codons, white box, phosphatase domain, light blue box, regions of re-codonised sequence (R.R.). (1C) Diagnostic PCR analysis of genomic DNA (gDNA) from a transgenic PP7 parasite line verifying successful modification of target loci by SLI to produce PP7-HA:loxP. Efficient excision of ’floxed’ sequences is observed upon treatment with RAP. Track C represents amplification of a control locus (PKAr) to check gDNA integrity. PCRs 1-4 are represented in the schematic locus in panel 1B. PCR 1 screens for the WT locus, PCR 2 for 5’ integration, PCR 3 for the excision of the ’floxed’ sequence and PCR 4 for 3’ integration. See Table 1 for sequences of all primers used for PCR. Sizes for expected amplification products are as follows: C, control locus (primers 5 and 6) 836 b.p. PCR 1 (primers 7 and 8) 1296 b.p, PCR2 (primers 7 and 9) 3559 b.p, PCR3 (primers 7 and 10) 1362 b.p. (RAP) and PCR 4 (primers 11 and 8) 1013 b.p. (1D) Western blot analysis of expression (DMSO, control) and ablation (RAP) of PP7-HA from highly synchronous mature schizonts. Expression of GAPDH (PF3D7_1462800) is shown as a loading control. (1E) IFA analysis showing the diffuse peripheral localisation of PP7-HA and the loss of expression upon RAP treatment (16 h post invasion). Over 99% of all RAP-treated PP7-HA:loxP schizonts examined by IFA showed diminished HA expression in three independent experiments. Scale bar, 2 µm.

**Table 1.**
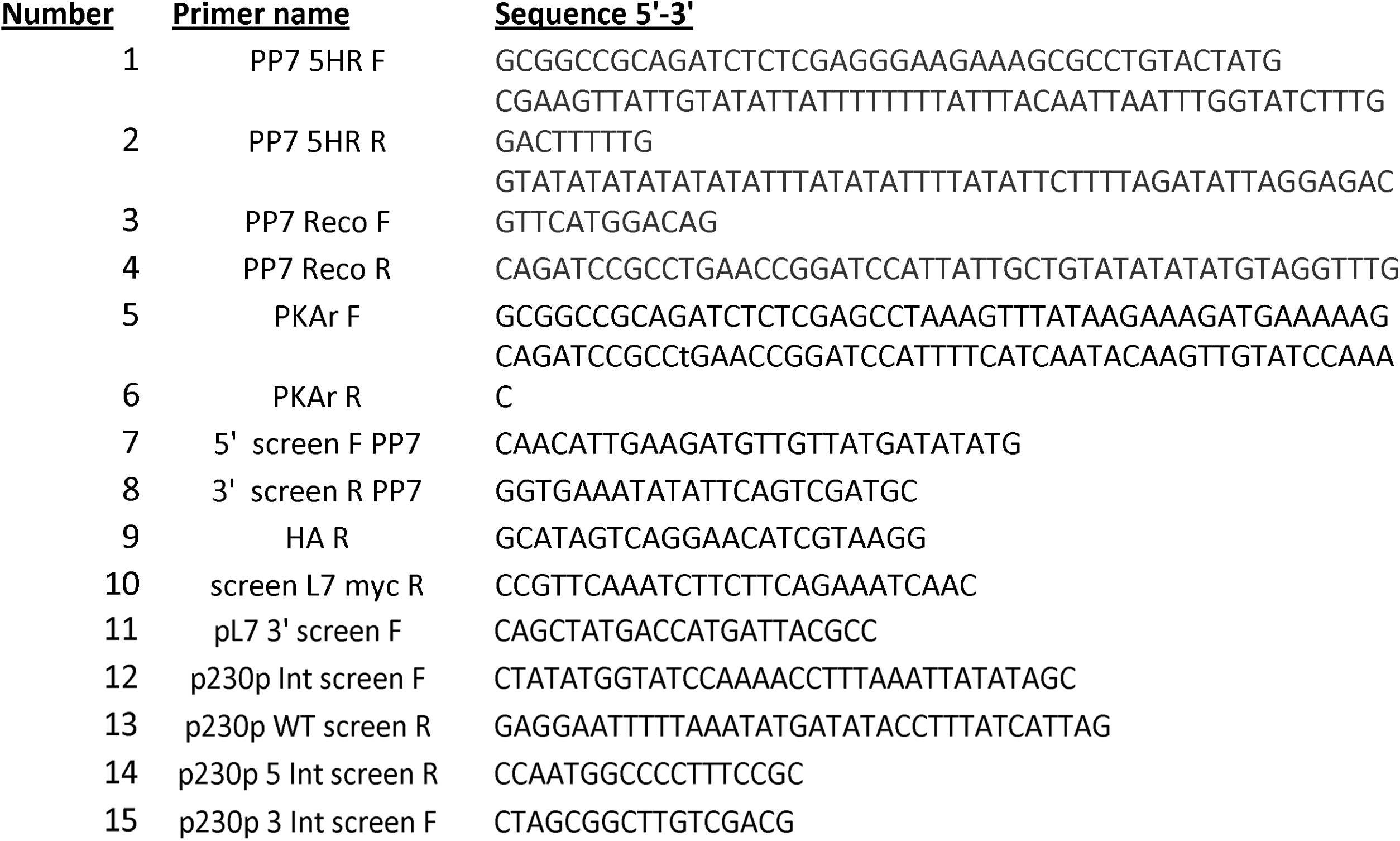
primers used in this study.

### *P. falciparum* PP7 is expressed in mature schizonts and is essential for asexual blood stage parasite growth

To investigate the biological function of PP7 in *P. falciparum* blood stages, we generated a transgenic parasite line (called PP7-HA:loxP) in a manner designed to allow determination of both the subcellular location of PP7 and the consequences of conditional disruption of PP7. The mutant line was generated on the genetic background of a 3D7 *P. falciparum* line that stably expresses dimerisable Cre (DiCre), the Cre-recombinase activity of which is induced in the presence of rapamycin (RAP) (23). The PP7 gene was ‘floxed’ such that treatment with RAP would lead to excision of DNA sequences encoding the phosphatase catalytic domain and all downstream features (Fig 1B). At the same time the gene was also modified by fusion to sequence encoding a C-terminal triple hemagglutinin (HA) epitope tag (Fig 1B).

Successful modification of the PP7 locus in PP7-HA:loxP parasites was verified by PCR (Fig 1B and C) and western blot, which also showed loss of expression following RAP-induced truncation of the tagged PP7-HA (Fig 1D). Immunofluorescence analysis (IFA) of the transgenic line revealed a partly peripheral signal in individual merozoites within mature segmented schizonts (Fig 1E).

To investigate the essentiality of PP7, highly synchronised young ring-stage cultures of PP7-HA:loxP were treated with RAP to induce excision of the sequence encoding the phosphatase catalytic domain at 4 hours (h) post invasion (Fig 1B). RAP-treated PP7-HA:loxP parasites seem to mature normally through the cycle of RAP-treatment (cycle 0). The RAP-treated schizonts produce the same number of merozoites (Fig S2A), have the same DNA content as schizonts treated with Vehicle (DMSO) only (Fig S2B), egress and re-invade normally (Fig S2C and D, movie S1). However, early in the next cycle (cycle 1), the parasites underwent severe growth arrest (Fig 2). Examination of the cycle 1 parasites by Giemsa-staining showed that the development of early ring stages was adversely affected in RAP-treated PP7-null parasites, with an accumulation of pycnotic and abnormal parasites that appeared immediately after invasion. In contrast, new rings of control PP7-HA:loxP parasites treated with DMSO, developed normally and went on to form schizonts (Fig 2 inset, Fig 5A and 5B). It was concluded that PP7 is essential for asexual blood stage survival and that it plays a role in formation of ring-stage parasites soon after erythrocyte invasion.

**Figure 2.**
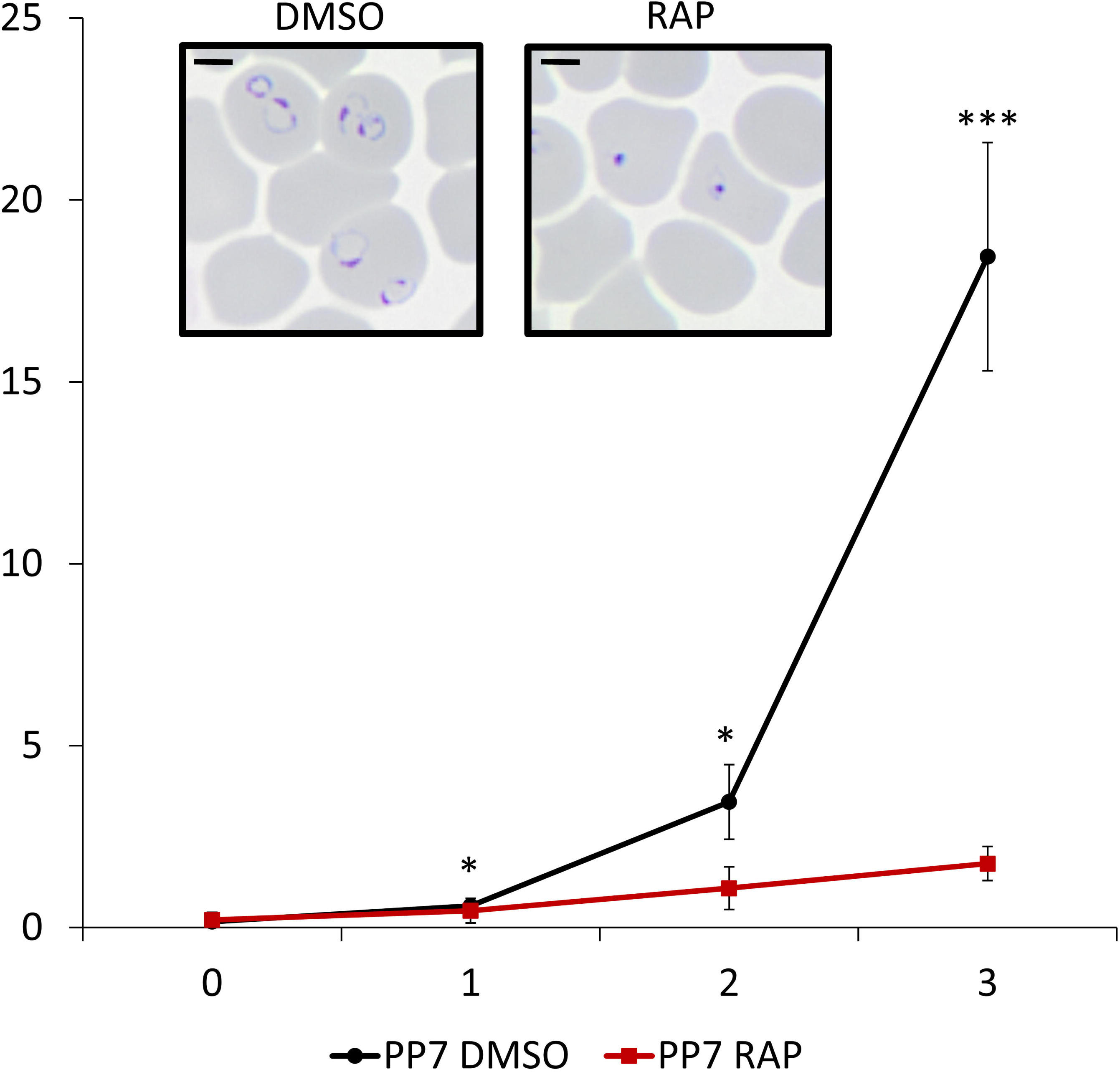
Growth curves showing parasitaemia as measured by flow cytometry of PP7-HA:loxP parasites treated with DMSO (vehicle only control, blue) or RAP (red). Means from three independent biological replicates are plotted. Error bars, SD. Inset, Giemsa-stained thin blood films showing ring-stage parasites following egress of synchronous DMSO- and RAP-treated PP7-HA:loxP schizonts. Ring formation occurs in DMSO-treated PP7-HA parasites, parasites did not develop beyond the early ring stage in RAP-treated parasites. Student’s t test for the comparison of conditions between DMSO and RAP treatments, * :P<0.05,*** :P<0.001.

### PP7 is a calcium-dependent phosphatase

Since the domain structure of *P. falciparum* PP7 includes putative domains involved in calcium sensing, we sought to investigate whether PP7 is a calcium-dependent phosphatase. For this, we isolated PP7 from extracts of PP7-HA:loxP parasites by immuno-precipitation with magnetic anti-HA conjugated beads (Fig 3A). The immobilized PP7 was then examined for phosphatase activity in the absence or presence of Ca^2+^ or chelating agents. As shown in in Fig 3B, PP7 displayed phosphatase activity which was strongly enhanced by the presence of Ca^2+^ and inhibited by calcium chelators (Fig 3B). These results indicate that parasite-derived PP7 is a functional phosphatase and that its activity is Ca^2+^ dependent.

**Figure 3.**
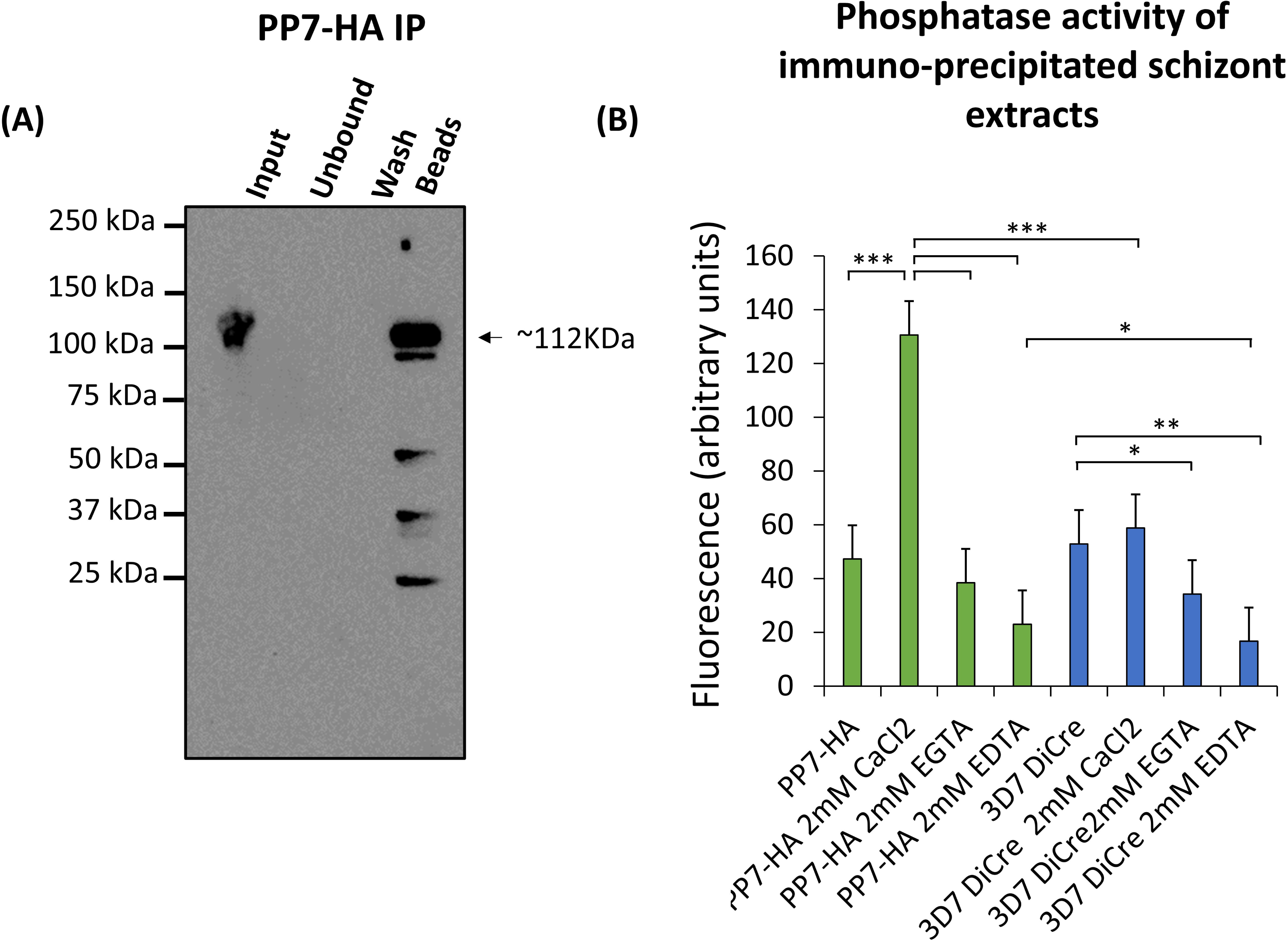
(3A) Western blot showing immunoprecipitation (IP) of PP7-HA from schizont extracts. Black arrow indicates the predicted mass of PP7-HA. Minor lower molecular weight degradation products are seen. Images are representative of two independent biological repeats. (3B) Phosphatase assay of immuno-precipitated PP7-HA conjugated beads. PP7 conjugated bead conditions are shown in green and control beads from 3D7 DiCre parental schizont lysate immuno-precipitation are displayed in blue. Results are averages of three independent biological repeats. Error bars, SD, * :P<0.05, ** :P<0.01, *** :P<0.001, Student’s t test for the comparison between conditions.

### PP7 interacts with calmodulin and CDPK1

The observed calcium-dependence of PP7 activity raised the possibility that it may play a role in calcium-mediated signalling. To evaluate this and to gain further mechanistic insight into the role of PP7, we sought to identify potential PP7-interacting partner proteins. For this, immuno-precipitated material from PP7-HA:loxP and 3D7 DiCre (negative control) schizonts were analysed by mass spectrometry to identify co-purifying protein species with and without the presence of additional Ca^2+^. As shown in Fig 4A and Supplementary Table 1, this analysis revealed that PP7 interacts with calmodulin and calcium-dependent protein kinase 1 (CDPK1), an enzyme involved in parasite invasion and development (9). This finding was confirmed by western blot analysis of PP7 immuno-precipitates using a CDPK1-specific antibody (Fig 4B). The mass spectrometric analysis also identified a conserved protein of unknown function (PF3D7_0423300) that was only enriched in the absence of added Ca^2+^, suggesting a calcium-dependent interaction of PP7 and PF3D7_0423300. Collectively, these results were interpreted as indicating a role for PP7 in calcium-mediated signalling in *P. falciparum*.

**Figure 4.**
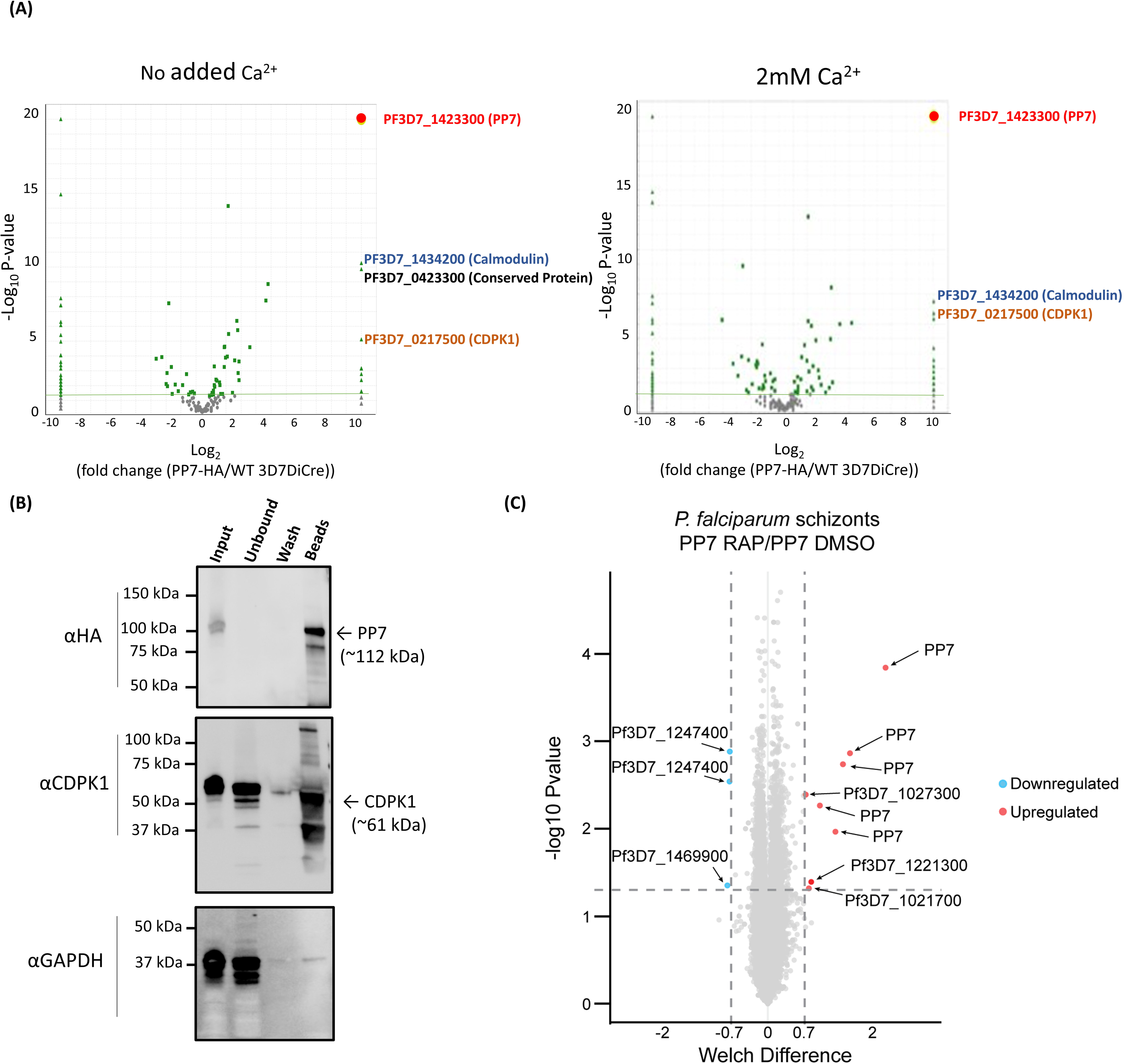
(4A) Mass spectrometric identification of interacting partners of PP7. The right panel shows immuno-precipitation with no added Ca^2+^ ions, left panel shows immuno-precipitation in the presence of additional 2 mM Ca^2+^ (present during detergent extraction, immuno-precipitation and washes). Volcano plot of P values versus the corresponding log2 fold change in abundance compared to 3D7DiCre control samples (Fischer’s exact test). Plotted by analysing proteins enriched through IP (panel 3A) by mass spectrometry. Green line indicates p=-2log10 and green dots represent peptides where p<-2log10. Peptides for PP7 were enriched to p<-19log10. (4B) Western blot analysis of immuno-precipitated PP7:HA-loxP schizont lysates with anti-HA beads, using the HA antibody (upper), the CDPK1 antibody (middle) and the GAPDH antibody (lower) (used as a negative control cytosolic marker to detect potential non-specific binding). (4C) Volcano plot showing the changes in detection of phospho-sites between DMSO- and RAP-treated PP7-HA:loxP. The negative log10 transform of the p-value–derived Welch-corrected t test comparing five DMSO- and five RAP-treated replicates is plotted against the log2-transformed fold change in reporter ion intensity (DMSO/RAP). Significance of p < 0.05 is denoted by the horizontal line. Vertical lines indicate –0.7 and 0.7 log2 fold change. Light blue circles correspond to significantly hypophosphorylated peptides and red circles to significantly hyperphosphorylated peptides.

### Loss of PP7 results in phosphoproteomic changes

To gain further insight into the mechanisms through which PP7 controls early ring-stage development, we profiled the PP7-dependent phosphoproteome from schizonts of PP7-null and control parasites cultures, reasoning that these are most likely to represent the developmental lifecycle stages in which Ca^2+^-dependent signalling could plausibly exert control over PP7 activity. Phosphopeptides enriched from trypsin-digested protein extracts of control and RAP-treated PP7-HA:loxP parasites were examined by tandem mass spectrometry using isobaric labelling for quantification. This showed that whilst phosphorylation sites of PP7 were significantly down-regulated in the PP7-null parasites, as expected, phosphosites of only four other proteins showed significant down- or up-regulation: the up-regulated sites were on the peptidyl-prolyl cis-trans isomerase FKBP35 (PF3D7_1247400) and on PfMGET (Pf3D7_1469900), whilst the down-regulated sites were on a putative EF hand domain-containing protein (PF3D7_1221300) which was found to be expressed in late schizonts in a transcriptomic study (24), on a putative peroxiredoxin (PfnPRx-Pf3D7_1027300) and on Pf3D7_1021700; a VPS13 domain-containing protein (Fig 4C, Table S2). Gene ontology (GO) enrichment analysis on hyper- and hypo-phosphorylated proteins suggested, primarily, deregulation in protein dephosphorylation and NADH dehydrogenase activity (Fig S3). These results confirmed that loss of PP7 results in a selective set of phosphoproteomic changes in the parasite that could underlie the ring development phenotype.

### PP7 is not required for invasion but is essential for early ring-stage development

To further investigate how loss of PP7 affects asexual parasite development, we quantified and characterized schizont and resultant ring formation over time in control and RAP-treated (PP7-null) PP7-HA:loxP parasites. This showed no effects of PP7 deletion on the apparent numbers of schizonts and subsequent ring stage formation, as determined by flow cytometry (Fig 5C). However, microscopic analysis of Giemsa-stained parasites 2 h post-invasion showed that newly intraerythrocytic PP7-null parasites did not develop post-invasion, instead displaying a predominantly dot-like morphology (Fig 5A, 5B and Fig 6A). To analyse this ring-stage development defect in more detail we modified the PP7-HA:loxP line to express a cytosolic mNeon-Green marker (7), resulting in the PP7-HA-cyto-mNeon:loxP line in which we could observe the morphology of the cell cytosol by green fluorescence in live cells. Imaging of control or RAP-treated PP7-HA-cyto-mNeon:loxP parasites that had additionally been stained with a membrane marker BODIPY Tr ceramide and DNA stain Hoechst 33342 showed that RAP-treated (PP7-null) parasites formed pycnotic structures that failed to expand, in contrast with the clearly-defined ameboid structures in control parasites (Fig 6B). Together these data indicate a crucial role for PP7 in early ring-stage ameboid development.

**Figure 5.**
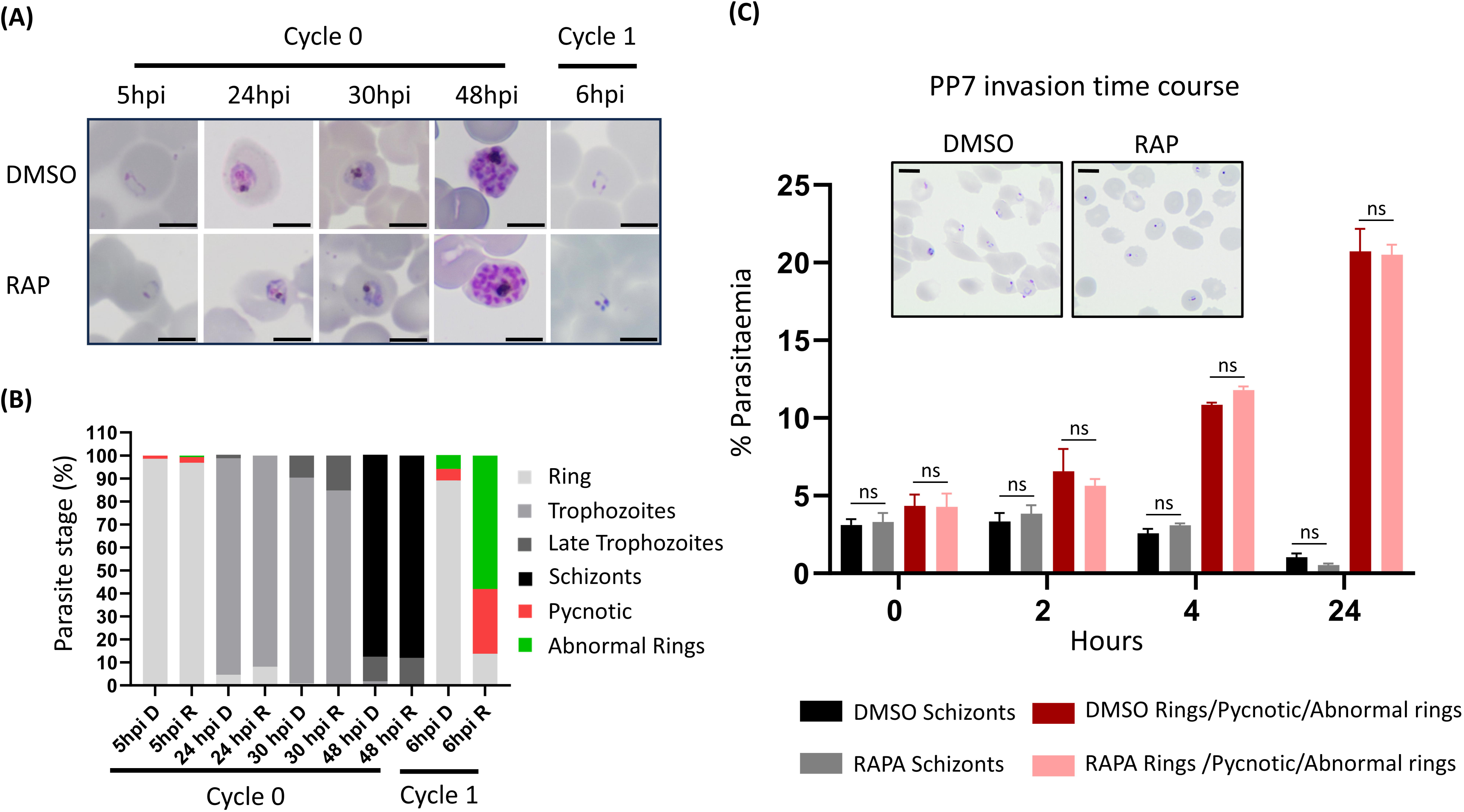
(5A) Giemsa-stained thin blood films showing development of PP7-HA:loxP parasites treated with DMSO or rapamycin during cycle 0 at 4 h post invasion for 2 h. Scale bars, 5 μm. (5B), Microscopic quantification of parasite developmental stages at each time point. Rapamycin-treated parasites displayed an accumulation of pycnotic and abnormal looking parasites at early life cycle stage. Counts are means of results of two independent experiments. Rapamycin treated parasites develop normally during the first cycle and are able to re-invade. Early on during the next cycle, an accumulation of abnormal rings and pycnotic forms appear, the parasites did not develop beyond this cycle. (5C) Flow cytometry analysis of ring formation in DMSO- and RAP-treated PP7-HA:loxP parasites treated in the previous cycle. Samples from highly-synchronized cultures treated at the ring stage the cycle prior to invasion and analysis, were taken in triplicate at the stated time intervals post invasion and stained with the DNA stain SYBR green. Samples were analyzed by flow cytometry, and the schizont and ring parasitemia values determined by gating high-signal and low-signal SYBR-positive cells, respectively. Mean parasitemia values (starting schizontemia adjusted to 2%) from two independent experiments are plotted. Error bars, SD; Student’s t test for the comparison of conditions between DMSO and RAP treatments (ns: nonsignificant). Insets are smears taken from cultures at 24h post invasion. Scale bars, 51μm.

**Figure 6.**
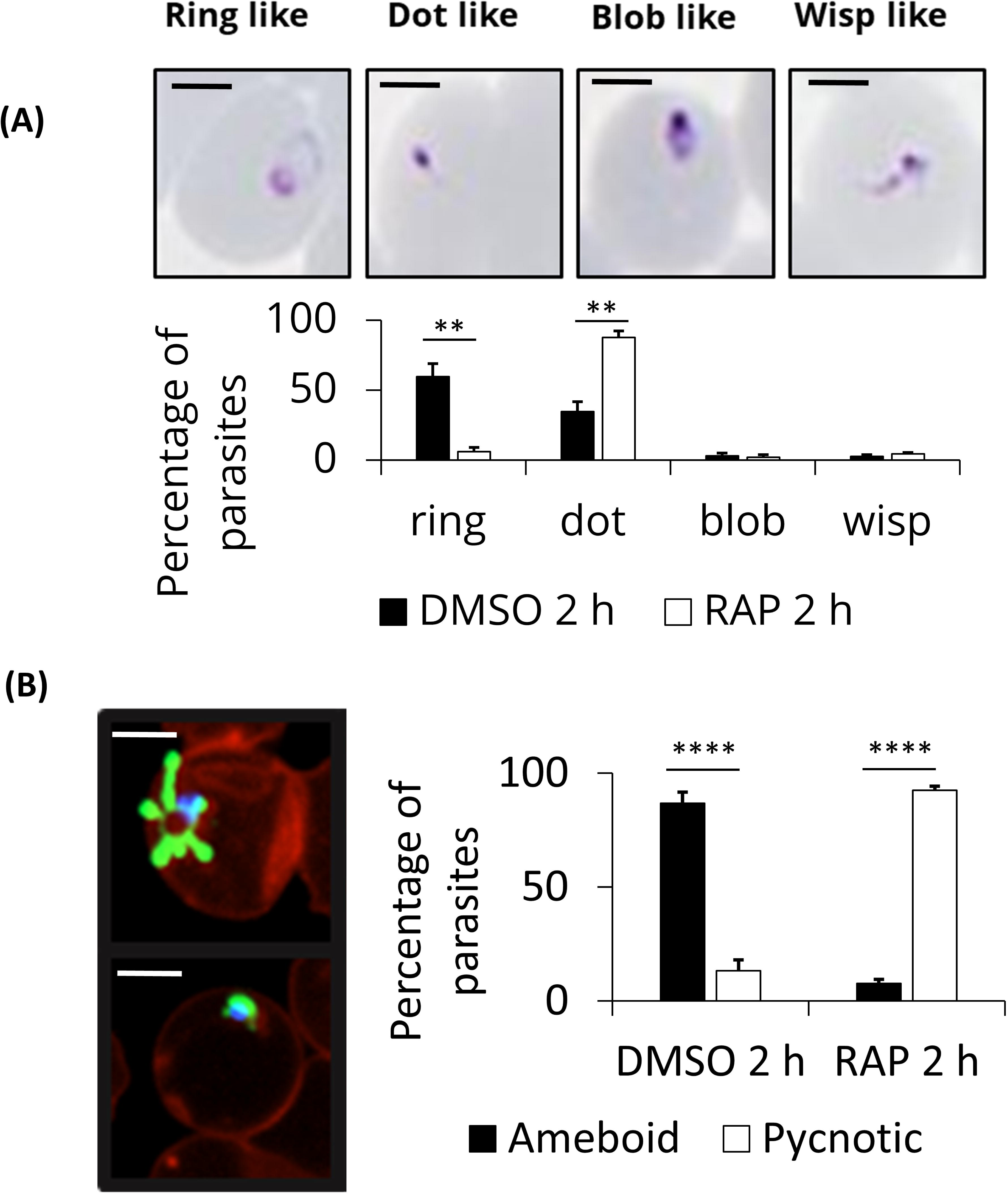
PP7 ablation results in arrest of ring-stage parasite development. (A) Tightly synchronized DMSO- or RAP-treated PP7-HA:loxP parasites were arrested with C2 block and subsequently washed and left to invade in fresh pre-warmed medium. After a further 2 h, thin blood smears were taken, and Giemsa stained. Ring-stage parasite morphology was classified by manual counting. A minimum of 100 cells were counted per replicate and scored for morphology, scale bar 2 µm. (B) Tightly synchronized DMSO- or RAP-treated PP7-HA-Cyto-mNeon:loxP parasites were arrested with C2 and subsequently washed and left to invade in fresh pre-warmed medium. Parasites were then stained with BODIPY TR ceramide and Hoechst 3342 to visualise membranes and nuclei respectively. Parasite samples were placed in an Ibid chamber slide and imaged by fluorescence microscopy. A minimum of 40 ring-stage cells were counted per condition and scored for ameboid or pycnotic morphology of parasites, scale bar 2 µm. Mean data are plotted for three independent experiments. Error bars, SD, ** :P<0.01, **** :P<0.0001, Student’s t test for the comparison of conditions between DMSO and RAP treatments.

## Discussion

The findings presented in this study shed light on the crucial role of PP7, a biochemically active serine/threonine protein phosphatase, in *P. falciparum* blood stages, expanding our understanding of the signalling machinery of this deadly parasite.

Our data unequivocally demonstrate that PP7 is indispensable for asexual blood stage survival in *P. falciparum*. Conditional disruption of PP7 led to a severe growth arrest, emphasizing its role in early asexual blood stage parasite replication. This aligns with previous findings from a global *P. berghei* knockout screen (4), reinforcing the notion that PP7 plays a conserved and vital role in *Plasmodium* species. While PP7 was found to be dispensable for erythrocyte invasion, its requirement for early ring-stage development is clear. The striking accumulation of pycnotic parasites in early ring stages upon PP7 disruption led us to investigate and confirm a role in amoeboid formation. This amoeboid form has recently been described (25) and remains an understudied aspect of early invaded parasite development. The developmental arrest observed in PP7-null rings may suggest a crucial role in the organization or maturation of structures critical for the development of newly invaded parasites that contribute to the amoeboid morphology. Interestingly, the *Toxoplasma* orthologue of PP7, *Tg*PP7, is also essential for parasite replication and virulence *in vivo* (26). In contrast to our findings with *P. falciparum* PP7, however, *Tg*PP7 was found to have a role in tachyzoite invasion. This implies the evolutionary co-option of PP7 to suit species-specific roles in apicomplexan invasion and early invaded parasite development. Further work is warranted to elucidate the precise mechanisms underlying these differences.

The discovery of PP7 as a calcium-dependent phosphatase is a significant revelation. The pronounced increase in PP7 activity in the presence of Ca^2+^, which could be reversed by calcium chelation, implies a role in calcium signalling, which is pivotal during various transitions during the asexual blood stage lifecycle, including erythrocyte invasion and egress (6, 9, 27, 28). Our results suggest that the calcium-dependent phosphatase activity of PP7 plays a crucial role in regulating key calcium-dependent events in the parasite life cycle.

These conclusions are supported by our identification of calmodulin and calcium-dependent protein kinase 1 (CDPK1) as interacting partners of PP7. Calmodulin is a known calcium sensor, and CDPK1 plays a crucial role in parasite invasion and development. The apparent interaction of PP7 with these proteins strongly suggests a role for PP7 in orchestrating crucial calcium-dependent signalling cascades. Given that PP7 is a high confidence PKAc substrate (7), it is plausible to suggest that PP7 may act as molecular player in the interaction between the PKAc and calcium-mediated cell signalling pathways in blood stage schizonts. Indeed, cross-talk between these two pathways has been previously observed (29, 30) but the precise molecular processes governing this remain to be elucidated. We suggest that this role could be fulfilled in part by PP7. (26)

The phosphoproteomic profiling of PP7-dependent processes in mature schizonts represents a rich dataset for further exploration. We detected five proteins other than PP7, of which the phosphorylation status is altered significantly upon loss of PP7. Among them, three (PF3D7_1021700, PF3D7_1027300, PF3D7_1221300) were also found to be deregulated in the PKAc null parasite phosphoproteome (7). The putative EF hand domain-containing protein PF3D7_1221300 is predicted to be non-essential (31), but nonetheless underscores the potential role of PP7 in calcium signaling. The peptidyl-prolyl cis-trans isomerase FKBP35 (PF3D7_1247400) is predicted to be essential (31) and is most highly expressed in the early schizont stage (24). FK506-binding proteins have been shown to catalyze the cis-trans isomerization of proline imidic peptide bonds (32) and have a role in protein folding (33) but little is known of the specific cellular roles in model organisms. The potential role of PP7 in regulation of protein folding processes presents an interesting avenue for future research. The only other protein predicted to be essential is Pf3D7_1021700 (31)and harbors a VPS13 domain. VPS13 proteins are known lipid transporters, targeted to distinct membranes and particularly membrane contact sites. In yeast, VPS13 is involved in membrane expansion(34). Although plausible that it has a similar role in *Plasmodium*, contributing to the ablation of ring formation, no mechanistic evidence is currently available. It will be interesting to investigate further whether PP7 disruption affects phosphorylation of PKAc substrates, especially in the presence of calcium signalling, which will imply a role in regulating PKAc activity. Given the importance of cAMP and calcium signalling in parasite biology, the modulation of PKAc or PKAr phosphorylation by PP7 is an exciting avenue for future research.

In conclusion, our findings underscore the multifaceted role of PP7 in *P. falciparum* asexual blood stage development. The essentiality of PP7, its calcium-dependent phosphatase activity, and its interactions with key signalling molecules, together with its role in some PKAc substrate phosphorylation in schizonts, emphasize the importance of PP7 in the cell biology of the parasite. Our work opens new avenues for research into the mechanisms underlying PP7 functions, its contribution to parasite survival, and its potential as a target for anti-malarial drug development.

## Acknowledgements

This work was also supported by funding from the Wellcome Trust to DAB (220318/Z/20/Z) and to MJB from the Wellcome Trust (220318/A/20/Z) and the Francis Crick Institute (https://www.crick.ac.uk/) which receives its core funding from Cancer Research UK (CC2129), the UK Medical Research Council (CC2129), and the Wellcome Trust (CC2129).

## Data availability

The mass spectrometry proteomics data have been deposited to the ProteomeXchange Consortium via the PRIDE (35) partner repository (http://proteomecentral.proteomexchange.org) with the dataset identifier PXD051277.

## Materials and Methods

### *P. falciparum* culture and synchronisation

*P. falciparum* erythrocytic stages were cultured in human erythrocytes (National Blood Transfusion Service, UK) and RPMI 1640 medium (Life Technologies) supplemented with 0.5% Albumax type II (Gibco), 50 μM hypoxanthine, and 2 mM L-glutamine at 37^0^C and supplied with (gas composition). Synchronous parasite cultures were obtained as described previously (36). Briefly, late segmented schizonts were enriched by centrifugation on a 63% Percoll (GE Healthcare) cushion, followed by the addition of fresh erythrocytes to allow invasion for 1–2 h with continuous shaking. Remaining schizonts were then lysed by sorbitol treatment to yield highly synchronous ring-stage cultures. In all cases, induction of DiCre activity when required was by treatment for 2–4 h with 100 nM rapamycin (RAP; Sigma) as described previously (37, 38). Control parasites were treated with vehicle only (0.1% v/v DMSO).

The PP7-HA:loxP line was generated from the DiCre-expressing 3D7 *P. falciparum* clone (23) using SLI (39) with a plasmid containing a SERA2loxPint followed by a triple-HA tag and an in frame *Thosea asigna* virus 2A (T2A) ribosomal skip peptide and NeoR cassette with a downstream loxP and PbDT 31UTR sequences as described previously (7). To ensure appropriate recombination to drive gene excision and C-terminal tagging, a re-codonised version of the C-terminal portion of PP7, containing the phosphatase catalytic domain and downstream EF hands, was synthesised commercially (IDT) and inserted downstream of the SERA2loxPint and upstream of the 3×HA tag (Figure S4). The sequence of the ∼800 bp upstream homology region is shown in Figure S4A.

Oligonucleotide primer sequences used in diagnostic PCR to detect integration and excision of transgenes, and the sequences of re-codonised regions, are provided below in Tables 1 and S4.

PP7-HA:loxP mNeon was generated by transfection of PP7-HA:loxP with a construct targeting the p230p locus. A linearised donor DNA fragment which inserted mNeon in-frame with a T2A peptide and BSD selection marker when integrated, and a pDC2-based p230p targeting Cas9 gRNA plasmid was co-transfected. Parasites were left to grow for two days post transfection followed by treatment with 5 µg/ml blasticidin (BSD) to select for integrants. After the emergence of BSD-resistant parasites gDNA was screened for correct integration.

### Parasite sample preparation and western blot

Parasite extracts were prepared from Percoll-purified schizonts treated with 0.15% w/v saponin to remove erythrocyte material. To solubilise parasite proteins, PBS-washed saponin-treated parasite pellets were resuspended in three volumes of NP-40 extraction buffer (10 mM Tris, 150 mM NaCl, 0.5 mM EDTA, 1% NP40, pH 7.5, with 1× protease inhibitors (Roche). Samples were gently vortexed and incubated on ice for 10 min followed by centrifugation at 12,000g for 10 min at 4°C. For western blot, SDS-solubilised proteins were electrophoresed on 4%-15% Mini-PROTEAN TGX Stain-Free Protein Gels (Bio-Rad) under reducing conditions and proteins transferred to nitrocellulose membranes using a semidry Trans-Blot Turbo Transfer System (Bio-Rad). Antibody reactions were carried out in 1% skimmed milk in PBS with 0.1% Tween-20 and washed in PBS with 0.1% Tween-20. Appropriate horseradish peroxide-conjugated secondary antibodies were used, and antibody-bound washed membranes were incubated with Clarity Western ECL substrate (Bio-Rad) and visualised using a ChemiDoc (Bio-Rad). Antibodies used for western blots presented in this work were as follows: anti-HA monoclonal antibody (mAb) 3F10 (diluted 1:2,000) (Roche); mouse anti-GAPDH mAb (1:20,000).

### Immunofluorescence assays

Thin blood films were fixed with 4% formaldehyde in PBS and permeabilised with PBS containing 0.1% (v/v) Triton X-100. Blocking and antibody binding was performed in PBS 3% BSA w/v at room temperature. Slides were mounted with ProLong Gold Antifade Mountant containing DAPI (Thermo Fisher Scientific). Images were acquired with a NIKON Eclipse Ti fluorescence microscope fitted with a Hamamatsu C11440 digital camera and overlaid in ICY bioimage analysis software. Antibodies used for IFA were as follows: anti-HA monoclonal antibody (mAb) 3F10 (diluted 1:100) (Roche); rabbit anti-GAP45 mAb (1:200).

### Flow cytometry

For growth assays, synchronous ring-stage parasites were adjusted to a 0.1% parasitaemia 1% haematocrit suspension and dispensed in triplicate into six-well plates. Triplicate samples of 100 μL were harvested at days 0, 2, 4 and 6 for each well and fixed with 4% formaldehyde 0.2% glutaraldehyde in PBS. Fixed samples were stained with SYBR green and analysed by flow cytometry.

For the measurement of egress and ring formation of highly synchronized DMSO- and RAP-treated PP7-HA:loxP parasites. A culture of PP7-HA:loxP with a 1 h invasion window was seeded in duplicate at 1% hematocrit at 2% parasitaemia and treated with DMSO or RAP at 4 h post invasion for 2 h. Triplicate 100 μL samples were taken and fixed with 4% formaldehyde 0.2% glutaraldehyde in PBS at hourly intervals from 45-53 h post invasion and at 69 h post invasion the subsequent day. Fixed samples were stained with SYBR green and analysed by flow cytometry. Schizont parasitaemia was determine by gating high signal SYBR positive cells. Ring parasitaemia was determined similarly but by gating low signal SYBR positive cells.

### Analysis of parasite development using Giemsa-stained samples

To analyse and measure the development of PP7-HA:loxP parasites, tightly synchronized schizonts were allowed to re-invade fresh erythrocytes. At 4 hours post invasion, the rings were treated with DMSO or rapamycin for 2 hours (cycle 0). At the indicated time point after invasion (cycle 0 and cycle 1), the parasites were pelleted, smeared on a microscope slides, fixed with methanol and stained with Giemsa. The parasites were then imaged and counted using an Olympus BX51 microscope equipped with an Olympus SC30 camera and a 100x oil objective, controlled by CellSens software.

### Merozoite number

The quantification of merozoites per schizont was conducted using Giemsa-stained smears of Percoll-enriched mature parasites. The cells were imaged on an Olympus BX51 microscope equipped with an Olympus SC30 camera and a 100x oil objective, controlled by cellSens software. Two independent experiments were conducted with a minimum of 15 schizonts counted per experiment. Statistical analysis was performed using GraphPad Prism v10.

### Parasite invasion rate

Parasite invasion assays were performed as described previously in (40). Briefly, parasites treated with either DMSO or rapamycin were diluted to a parasitaemia of 1% and a haematocrit of 2%. A 50 µl starting sample (designated as H0) was collected and fixed (PBS solution containing 8% paraformaldehyde, 0.01% glutaraldehyde). After a 4-hour incubation period, a second sample was collected and fixed (designated as H4). The parasites were labelled with SYBR Green I (1:5000, Life Technologies). The parasitaemia was quantified using an Attune cytometer (Thermofisher). Parasite invasion rate was determined as the ratio of parasitaemia at H4 to H0. Each experiment was conducted in triplicate, with a minimum of three biological replicates. Statistical analysis was performed using GraphPad Prism v10.

### Schizont DNA content analysis

Highly synchronous schizonts were fixed and labelled with SYBR Green I (1:5000, Life Technologies). The fluorescence of SYBR Green was analysed using an Attune cytometer (Thermofisher) with the following laser settings: forward scatter at 125 V, side scatter at 350V, and blue laser (BL1) 530:30 at 280V. For each sample, at least 100,000 cells were analysed with three biological replicates performed in triplicate. Data analysis was carried out using FlowJo software.

### Parasite egress assay and time-lapse and live fluorescence microscopy

Highly synchronous schizonts were isolated using Percoll centrifugation and incubated for 4 hours in medium containing the PKG inhibitor C2 to arrest egress (1 μM, (38). DMSO-treated parasites were treated with Hoechst to stain the nuclei and mixed with the unstained RAP-treated parasites. Prior to analysis, the schizonts were resuspended without C2 in fresh serum-free RPMI at 37°C to allow egress and immediately loaded onto a poly-L-lysine coated μ-slide VI 0.4 (Ibidi). The slides were transferred to a Nikon Ti-E inverted microscope housed in a pre-warmed 37 °C chamber with an atmosphere of 5% CO_2_.

Egress was imaged using a 63× oil immersion objective and an ORCA-Flash 4.0 CMOS camera (Hamamatsu). DIC (differential inference contrast) images were taken at a rate of 1 frame/10 sec for 30 minutes, fluorescence images were taken every 2 min to avoid bleaching. Videos were acquired and processed using Nikon NIS-Elements and Image J software.

### Analysis of PP7 rings stage morphology

A culture of PP7-HA:loxP-mNeonGreen parasites with 1 h synchronicity was split and treated with DMSO or RAP. Cultures were held pre-egress with 1 µM PKG inhibitor (C2) until 50 h post invasion at which point C2 was removed by washing with pre-warmed complete medium. Cultures were left to invade new host cells for 1 h with continuous shaking and staining with BODIPY-Tr ceramide (Thermo). Subsequently a sample of each culture was taken. For Giemsa-stained blood film analysis samples were smeared and fixed with methanol followed by Giemsa staining. For fluorescence microscopy, samples were washed and diluted to 0.1% haematocrit, and placed into a chamber of a poly-L lysine coated Ibidi chamber slide to adhere for 10 minutes at 37 degrees. Adherent cells were then imaged on a NIKON Eclipse Ti fluorescence microscope fitted with a Hamamatsu C11440 digital camera and overlaid in ICY bioimage analysis software.

### Immuno-precipitation

Tightly synchronised schizonts (∼45 h post invasion) of PP7-HA:loxP and 3D7DiCre parental parasites were enriched on a 70% Percoll cushion. The schizonts were treated for 3 h with 1 μM C2 (to arrest egress) after which the cultures were treated with 0.15% saponin in PBS containing cOmplete Mini EDTA-free Protease and PhosSTOP Phosphatase inhibitor cocktails (both Roche) for 10 min at 4°C to lyse the host erythrocytes. Samples were washed twice in PBS containing protease and phosphatase inhibitors, snap-frozen and pellets stored at −80°C. Parasite pellets (70-100 μl packed volume) were resuspended in three volumes of NP-40 extraction buffer (10 mM Tris, 150 mM NaCl, 0.5 mM EDTA, 1% NP40, pH 7.5, with 1× protease inhibitors (Roche). Samples were gently vortexed and incubated on ice for 10 min followed by centrifugation at 12,000g for 10 min at 4°C. Clarified lysates were then added to anti-HA antibody-conjugated magnetic beads (Thermo Scientific) which had been equilibrated in NP-40 extraction buffer. Samples were incubated at room temperature for 2 h on a rotating wheel after which beads were precipitated using a magnetic sample rack. The supernatant was removed, and beads washed three times with NP-40 extraction buffer followed by three washes with extraction buffer lacking detergent.

### Mass spectrometry of immuno-precipitated material

Washed beads were resuspended in trypsinisation buffer (50 mM ammonium bicarbonate, 40 mM 2-chloroacetamide and 10 mM Tris-(2-carboxyethyl) phosphine hydrochloride) and samples reduced and alkylated by heated to 70°C for 5 minutes. 250 ng of trypsin was added to the samples and heated at 37°C overnight with gentle agitation followed by filtration using a 0.22 µm Costar® Spin-X® centrifuge tube filter (Sigma). Samples were then run on a LTQ-Orbitrap-Velos mass spectrometer (Thermo Scientific). Search engines, Mascot (http://www.matrixscience.com/) and MaxQuant (https://www.maxquant.org/) were used for mass spectrometry data analysis. The PlasmoDB database was used for protein annotation. Peptide and proteins having minimum threshold of 95% were used for further proteomic analysis and peptide traces analysed using Scaffold5. Enrichment was determined by comparing results from tagged lines with those of immuno-precipitated material from 3D7DiCre parental parasites.

### Phosphatase assay

Assays were carried out using the EnzChek Phosphatase Assay Kit (Therno) according to the manufacturer’s procedures. Briefly, percoll purified schizont pellets of 50 µl were prepared for PP7-HA:loxP or 3D7DiCre parental parasites. Samples were processed by the methodology for immuno-precipitation. PP7-HA conjugated beads were then resuspended in a total volume of 650 µl of assay buffer (20 mM Tris, 150 mM NaCl, pH 7.5,). 50 µl of suspended bead slurry was dispensed in triplicate into wells of a 96-well plate for each test condition. 50 μl of reaction buffer containing additives for the conditions tested and difluoro-4-methylumbelliferyl phosphate (DiFMUP) substrate were added, to give a final concentration of 100 µM DiFMUP. A set of control wells were made with the addition of reaction buffer lacking DiFMUP. Plates were incubated at 37°C for 1 h and then read with Ex/Em of 358/455 nm on a spectramax i5 plate reader (Molecular Devices). Averaged arbitrary fluorescence for each condition was calculated by normalising to the negative control wells.

### Phosphoproteomics

The phosphoproteomics data presented are from an isobaric labelling experiment. Tightly synchronised, ring-stage PP7-HA:loxP were treated with 100 nM RAP or vehicle only (DMSO) and schizonts (about 45 h old) enriched on an approximately 70% Percoll cushion. The schizonts were treated for 2 h with 1 μM C2 (to arrest egress) and then washed to allow egress for 25 min, after which the cultures were treated with 0.15% saponin in PBS containing cOmplete Mini EDTA-free Protease and PhosSTOP Phosphatase inhibitor cocktails (both Roche) for 10 min at 4°C to lyse the host erythrocytes. Samples were washed twice in PBS containing protease and phosphatase inhibitors, snap-frozen in liquid nitrogen, and pellets stored at −80°C. Parasite pellets were resuspended in 1 mL 8 M urea in 50 mM HEPES, pH 8.5, containing protease and phosphatase inhibitors and 100 U/mL benzonase (Sigma). Proteins were extracted from the pellets using three 15 s bursts with a probe sonicator followed by a 10 min incubation on ice and a 30 min centrifugation at 14,000 rpm at 4°C. Protein content was estimated by a BCA protein assay and a 200 μg aliquot of each sample was taken for further processing. Samples were reduced with 10 mM dithiothreitol for 25 min at 56°C and then alkylated with 20 mM iodoacetamide for 30 min at room temperature. The alkylation reaction was quenched with an additional 10 mM dithiothreitol, and then each sample was diluted with 50 mM HEPES to reduce the urea concentration to <2 M prior to digestion. Proteolytic digestion was carried out by the addition of 4 μg LysC (WAKO) and incubated at 37°C for 2.5 h followed by the addition of 10 μg trypsin (Pierce) and overnight incubation at 37°C. After acidification, C18 MacroSpin columns (Nest Group) were used to clean up the digested peptide solutions and the eluted peptides dried by vacuum centrifugation. Samples were resuspended in 50 mM HEPES and labelled using the 0.8 mg Tandem Mass Tag 10plex isobaric reagent kit (Thermo Scientific) resuspended in acetonitrile. Labelling reactions were quenched with hydroxylamine, and a pool was made of each set of samples. Acetonitrile content was removed from the pooled TMT solution by vacuum centrifugation and then acidified before using a Sep-Pak C18 (Waters) to clean up the labelled peptide pool prior to phosphopeptide enrichment. The eluted TMT-labelled peptides were dried by vacuum centrifugation and phosphopeptide enrichment was subsequently carried out using the sequential metal oxide affinity chromatography (SMOAC) strategy with High Select TiO2 and Fe-NTA enrichment kits (Thermo Scientific). Eluates were combined prior to fractionation with the Pierce High pH Reversed-Phase Peptide Fractionation kit (Thermo Scientific). The dried TMT-labelled phosphopeptide fractions generated were resuspended in 0.1% TFA for LC-MS/MS analysis using a U3000 RSLCnano system (Thermo Scientific) interfaced with an Orbitrap Fusion Lumos (Thermo Scientific). Each peptide fraction was pre-concentrated on an Acclaim PepMap 100 trapping column before separation on a 50-cm, 75-μm I.D. EASY-Spray Pepmap column over a 3-h gradient run at 40°C, eluted directly into the mass spectrometer. The instrument was run in data-dependent acquisition mode with the most abundant peptides selected for MS/MS fragmentation. Two replicate injections were made for each fraction with different fragmentation methods based on the MS2 HCD and MSA SPS MS3 strategies described. The acquired raw mass spectrometric data were processed in MaxQuant (version 1.6.2.10) for peptide and protein identification; the database search was performed using the Andromeda search engine against the Homo sapiens canonical sequences from UniProtKB (release 2018_05) and *P. falciparum* 3D7 sequences from PlasmoDB (18). Fixed modifications were set as Carbamidomethyl (C) and variable modifications set as Oxidation (M) and Phospho (STY). The estimated false discovery rate was set to 1% at the peptide, protein, and site levels. A maximum of two missed cleavages were allowed. Reporter ion MS2 or Reporter ion MS3 was appropriately selected for each raw file. Other parameters were used as preset in the software. The MaxQuant output file PhosphoSTY Sites.txt, an FDR-controlled site-based table compiled by MaxQuant from the relevant information about the identified peptides, was imported into Perseus (v1.4.0.2) for data evaluation.

For a phosphorylation site to be considered regulated, the following cut-offs were applied: P-value <10.05, Welch difference >10.7 or < -0.7 and localisation probability >0.7. Significantly hyper- and hypo-phosphorylated proteins were used for GO enrichment analysis. GO IDs were extracted from the *P. falciparum* annotation file (PlasmoDB.org). Enriched GO terms were identified as described before (PMID: 25011111). Plots were made using R package ggplot2 (https://ggplot2.tidyverse.org).

