## Supplementary material for "*Plasmodium falciparum* protein phosphatase PP7 is required for early ring-stage development"

**Supplementary Figure S1**

(A)

MENYNIEDVVMIYGHKAIIESIQNVENVKSEIQPNKIYVGKYVNKKGYSDGTYRKKKIYTPQNDELVILCSYNSIKPY**NE**

**EESACTMIQKMFRGYQGRKSFH**SFVCCTVWRKFDHIHEYITLNNHDEIYKPLIKTIKRDIKKGIIQFPSKCFSSNSNQFS

RVSTTSYESSNYNENV**PRLKDKIDRTFATEMFHFFLTSKEIILPYNLVHKVLIKTKKMLEENIKSSVINLDMSKKSKDTK**

**LIILGDVHGQLHDVLWLFNRFGIPSSTNIYIFNGDIADRGENATEIFILLFIFKLSCNDSVIINRGNHECSYMNEVYGFY**

**NEVLSKYDNTIFDLFQNIFELLGLAVNVQNQIFVVHGGLSRYQDITLKEIDELDRKKQEILHPEQYEDIVIFDLLWSDPQ**

**KKEGIGGNARGNNCITFGPDVTEMFLKNNNLDILIRSHQVPKTLKGIESHHEGKCITLFSASNYCNKIKNLGAAIIFNQD**

LTFEVQEYMSPSLEVIRETFEENQKLREKVLHCSKIVELEKNEQKNNNKLSTEGLMNDIINCLSTIICNEKNSLWNNLYK

QDKDKKGVVHINIWKEELGKLSKAKKVPWIYLCRKLKMIEDYHVNYNNILSRFKINYAPNEKFLNTEWKNECFEHLYEAL

LKADLS**LRETLMVFDKNLDGKVSFAEFEQVLRDLNIDLSNEQIRILVRLINSNSLCNNTNLQENDKIDVAEFI**GKMRVCY

RLSINKDYVNNEKIQKLIETIGKHILSDSADTANYHYKFYEENNERHNSERRKRSSVIKSV**ALFQKFKNYDNFGNGYLDY**

**NDFVKAIKNFDMNKISKEVEFEVDDDILMELAKSIDITKSSKINFLEFLQAFY**VVNKSKYSYVDEIWCHICTVIYENKVA

LKKCMKYLEDTMQGKITSIHFRYILLELNKILQEHNFEMNKPLTDEQIDLLAYTVETDDKVDYVEFFNSFKPTYIYSNN

(B)

```
PfPP7     IKPYNEEESACTMIQKMFRGYQGRKSFHSFVCCTVWRKFDHI  116
PvPP7     IRPFSETESASIRIQRAFRGYRARKEFHSFVSSLVWRKFEHL  116
TgPP7     ARPYDEKDCSATQLQAAWRSYRARQAFQDAAAFNFWNELDAF  118
DmrdgC    PPDLETTIRAAIFIQKWYRRHQARREMQRRCNWQIFQNLEYA   56
HsPPEF1   TRRSDTSLRAALIIQNWYRGYKARLKARQHYALTIFQSIEYA   51
              .    :.  :*  :* ::.*   :      .:..::

```

(C)

```
PfPP7     SKCFSSNSNQFSRV----STTSYESSNYNENVPRLKDKIDRTFATEMFHFFLTSKEIILP	204
PvPP7     SKCFSSNSNQFSRI----STTSYESTAYDENVPKLKDKIDKAFAMEVFHFFLTTKNYVLP	204
TgPP7     SRRRSSTRRSFGRSSVDASSISAENIPDDYAGPRVKWPITREFCLDLVDHYRDNPDVPLP	238
DmrdgC    GSAHVSVLDDKDDL----VE-----EFGDIVNAKIELPIRKNHIDLLIDVFRKKRGNRLH	137
HsPPEF1   NQSLESEQDMRDRW----DYVDSIDVPDSYNGPRLQFPLTCTDIDLLLEAFKEQQ--ILH	135
          .    *     .                .    :::  :       :.. :       *

```

**D** **HGQ** **D**	**G**

```
PfPP7     YNLVHKVLIKTKKMLEENIKSSVINLDMSKKSKDTKLIILGDVHGQLHDVLWLFNRFGIP	264
PvPP7     NNLVCKILTKTQKMLEENIKSSVIHVDMTKKSKDTKLIILGDVHGQLNDVLWLFNRFGLP	264
TgPP7     RKYALELIMAIADHYRETMKGAVVEVEIPKKG-GSRLVLVGDTHGQLNDVLWIFYKFGPP	297
DmrdgC    PKYVALILREAAKSLKQLPNISPVSTAVSQQ-----VTVCGDLHGKLDDLLVVLHKNGLP	192
HsPPEF1   AHYVLEVLFETKKVLKQMPNFTHIQTSPSKE-----VTICGDLHGKLDDLFLIFYKNGLP	190
           : .  ::    .  .:  : : :     ::     : : ** **:*.*:: :: : * *

```

**F** **GD** V**DRG** S	**K**	**RGNHE	F E**
PfPP7	**S**ST**N**I**Y**I**FNGD**IA**DRG**ENAT**EI**FIL**L**FIFK**L**SCNDS**V**II**NRGNHE**CSY**MN**EV**YGF**YN**E**VL	324

```
PvPP7     SSNNIYIFNGDIADRGENATEIFLLLFTFKLSNYDCVIINRGNHECSYMNEVYGFYNEVL	324
TgPP7     SATNVYLFNGDIADRGRYAVEIFMMLFAFKLQCPSSVVINRGNHESADMNEVYGFAQEVR	357
DmrdgC    SSSNPYVFNGDFVDRGKRGLEVLLLLLSLYLAFPNAVFLNRGNHEDSVMNARYGFIREVE	252
HsPPEF1   SERNPYVFNGDFVDRGKNSIEILMILCVSFLVYPNDLHLNRGNHEDFMMNLRYGFTKEIL	250
          *  * *:****:.***. . *::::*    *   . : :******   **  *** .*:
```

**F**	**HGG**	**R**	P

```
PfPP7     SKY---DNTIFDLFQNIFELLGLAVNVQNQIFVVHGGLSRYQDITLKEIDELDRKKQEI-	380
PvPP7     SKY---DSAVFDLFQGVFELLGLAVNIQNQIFVVHGGLSRYQDLTMREIDELDRKKHEI-	380
TgPP7     QKY---GGFMYQKFQEVFHLLPLCVVMEKRVFVVHGGLCRKDNVTLQHIDRLNRQRPCP-	413
DmrdgC    SKYPRNHKRILAFIDEVYRWLPLGSVLNSRVLIVHGGFSD--STSLDLIKSIDRGKYVSI	310
HsPPEF1   HKYKLHGKRILQILEEFYAWLPIGTIVDNEILVIHGGISE--TTDLNLLHRVERNKMKSV	308
           **      :   :: .:  * :   ::..::::***:.      :  :. ::* :

                                                       G      W D
```

PfPP7	-------------------------------------LHPEQY**E**DIV**IFD**L**LWSDPQ**KKE	403
PvPP7	-------------------------------------LHPEKY**E**DTI**IFD**L**LWSDPQ**KKN	403

```
TgPP7     -------------------------------------ACPHSFEDTLMFDLLWSDPQHES	436
DmrdgC    LRPPLTD--------------------------G---EPLDKTEWQQIFDIMWSDPQATM	341
HsPPEF1   LIPPTETNRDHDTDSKHNKVGVTFNAHGRIKTNGSPTEHLTEHEWEQIIDILWSDPRGKN	368
                                                   . *   ::*::****:
```

**RG**	F	**R H**	**G**	**T** **SA**P
PfPP7	**G**IGG**N**A-**RG**NN**C**IT**FGPD**V**T**EM**FL**KNNN**L**DIL**IRSH**QVPKTLK**G**I**E**SH**H**E**GK**CI**T**L**FSAS**	463
*PvPP7*	**G**IGG**N**A-**RG**NN**C**IT**FGPD**I**T**DL**FL**KKNNFDIL**IRSH**QVPKTLK**G**I**E**SH**H**E**GK**CI**T**L**FSAS**	462

```
TgPP7     GRGWST-RGADCIAFGPDITDAFLTKNNLEVCIRSHQVPTNLRGFEPVHDGRCVTLFSAS	495
DmrdgC    GCVPNTLRGAGVW-FGPDVTDNFLQRHRLSYVIRSHECKP--NGHEFMHDNKIITIFSAS   398
HsPPEF1   GCFPNTCRGGGCY-FGPDVTSKILNKYQLKMLIRSHECKP--EGYEICHDGKVVTIFSAS	425
          *   .: **     ****:*. :* . .:.  ****:     .* *  *:.: :*:****
```

**NY	N**
PfPP7	**NY**CNKIK**N**L**GA**A**I**IFNQD**L**TFEVQE**Y**MS 490
*PfPP7*	**NY**CNKIK**N**L**GA**A**I**VFNQD**L**TFEVQE**Y**MS 490

```
TgPP7     NYCGTTGNFGGVIIFEANLSFEIQEYMA  525
DmrdgC    NYYAIGSNKGAYIRLNNQLMPHFVQYIS  426
HsPPEF1   NYYEEGSNRGAYIKLCSGTTPRFFQYQV  453
          **     * *. * :      .. :*

```

(D)

```
PfPP7     ADLSLRETLMVFDKNLDGKVSFAEFEQVLRDLNIDLSNEQIRILVRLINSNSLCNNTNLQ	702
PvPP7     ADLSLRETLMVFDKNLDGKVSFAEFEQVLKDLNIDLSNEQIRILVRLINSNSLCNQDSSQ	702
TgPP7     ADLSLKETLMLFDRNCDGTVSFREFNELITELDVGLSEPQVRILMRLITASPAFNA----	729
DmrdgC    ------------------------------------------------------------	529
HsPPEF1   ------------------------------------------------------------	555

```

PfPP7	ENDKID**V**A**E**FIGKMRVCYRLSINKDYVNNEKIQKLIETIGKHILSDSADTANYHYK-FYE	761
PvPP7	ESDKID**V**A**E**FIGKMRVCYRLAINKEYTNNEKIRKLIETIGKHILADSADTANFHYR-FYE	761

```
TgPP7     AAGSIDVAEFLGRFRVVYSHVINDDKRSVPWLQRALHCIGKAILADKAEAANRHYEQQRQ	789
DmrdgC    EADGMSVMDALYAN----------------------------------------------	543
HsPPEF1   AH--STLVETLYRY----------------------------------------------	567
```

: : :

PfPP7	ENNERHNSERRKRSSVIKSVA**L**FQK**F**KNY**D**NFGN**G**YLDYND**F**VKA**IK**NFDMNKISKEVEF	821
PvPP7	ENNESAGSERRKRSSVIKSVA**L**FQK**F**KNY**D**NFGN**G**YLDYDD**F**VRA**IK**NFDMNKISKEVEF	821

```
TgPP7     DGNLGPDLNTRRRSSAVRAVALFQKFKDYNEGGDGYLSYADFVTGIKRLTIDE--EELGF	847
DmrdgC    ------------------KASLVAIFNIIDADNSGEITLDEFETAIDLLVAHM-----PG	580
HsPPEF1   ------------------RSDLEIIFNAIDTDHSGLISVEEFRAMWKLFSSHY-----NV	604
```

*	*: : .* : :*	. : .

PfPP7	EVD**D**DILME**LA**KSI**D**ITKSSK**I**NFL**EFL**Q**AF**Y**VV**N-KSKYSYVDEIWCHICTVIYEN**K**VA	880
PvPP7	DVD**D**EILME**LA**KSI**D**ITKSSK**I**NFL**EFL**Q**AF**Y**VV**N-KSKYSYVDEIWCHICTVIYEN**K**VA	880

```
TgPP7     ALTDEHLYKVAEAVDTTGSKRINYLEFLQAFHVVDSNSNNSAAEELWGQICTAIFQHKSS	907
DmrdgC    AYSKAEMLEKCRMMDLNGDGKVDLNEFLEAFRLSDLHRKEQQDENIRRRSTGRPSVAKTA	640
HsPPEF1   HIDDSQVNKLANIMDLNKDGSIDFNEFLKAFYVVHRYE-----------DLMKPDVTNLG	653
             .  : : .. :* . .  ::  ***:** : .                      : .
```

**Domain architecture and similarity of *P. falciparum* PP7 to EF-hand-containing protein phosphatases of higher eukaryotes**

Although the name PP7 was coincidentally attributed to phosphatases from both animals and plants, it has been suggested that they are not orthologues and that the PP7 nomenclature is best used for the plant phosphatases of this PPP sub-family and PPEF for the animal phosphatases(Andreeva and Kutuzov, 2009). PPEFs also have an IQ-type calmodulin-binding domain upstream of the catalytic domain. Plant PP7s have a different calmodulin-binding domain that forms an insert within the catalytic domain (Kerk et al., 2021).

(Supp 1A) Amino acid sequence of the *P. falciparum* PP7 (PF3D7_1423300) showing its conserved domains. The IQ calmodulin-binding domain is shown in red, the phosphoprotein phosphatase (PPP) catalytic domain in blue and the EF-hand domains in green. These coloured sequences are boxed in panels B-D.

(Supp 1B) A Clustal alignment of the IQ (calmodulin-binding) domain of *P. falciparum* PP7 with the *P. vivax* and *T. gondii* orthologues, and also with two protein phosphatases of higher eukaryotes which have the same domain architecture: the *Drosophila melanogaster* retinal degradation serine/threonine phosphatase rdgC and its human orthologue PPEF1. Residues shown in bold red are identical in all five sequences, those in bold black are identical in four of the five sequences. The Clustal assignment of identical and conserved residues between the five sequences are shown below the alignment.

(Supp 1C) A Clustal alignment, with minor manual modification, of the same five proteins showing the protein phosphatase catalytic domain. Residues shown in bold blue are identical in all five sequences, those in bold black are identical in four of the five sequences. Of the 44 residues conserved across PPP families, as defined previously (Barton et al., 1994), 38 are present in PfPP7 and are shown in bold red above the alignment, and the six not conserved are shown in black. Motifs that are highly conserved amongst all eukaryotic PPP motifs, as previously depicted, (Kerk et al., 2021) are boxed with the positions of metal-binding residues highlighted in green and those involved in catalysis and arginine residues that interact with the substrate, highlighted in yellow). The Clustal assignment of identical and conserved residues between the five sequences are shown below the alignment. The phosphatase catalytic domain of PfPP7 is boxed and corresponds to the sequence coloured blue in panel A.

(Supp 1D) A Clustal alignment of the same five proteins showing the EF-hand domains. Residues shown in bold green are identical in all five sequences, those in bold black are identical in four of the five sequences. The Clustal assignment of identical and conserved residues between the five sequences are shown below the alignment.

**Supplemental Figure 2:**

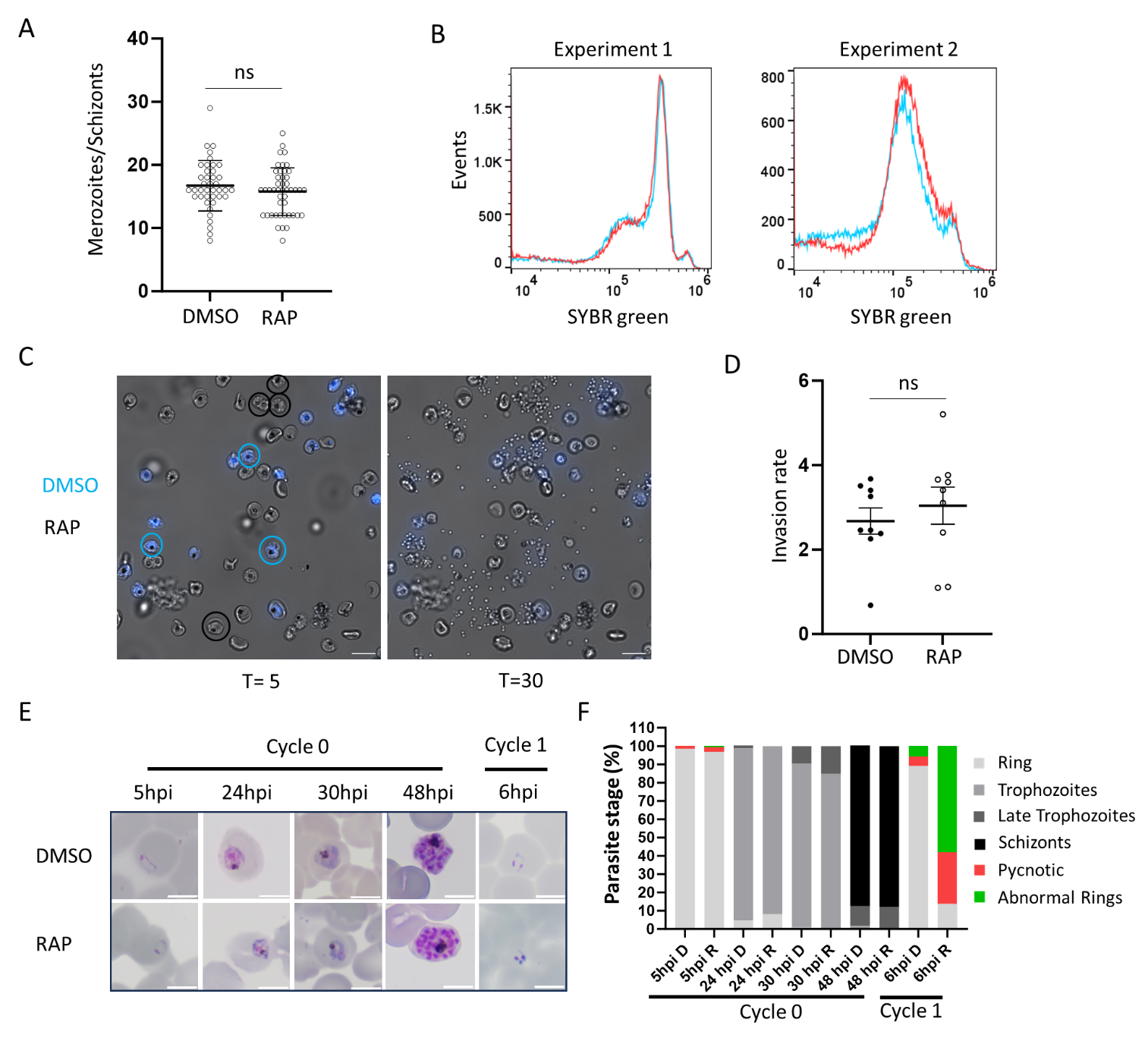

(Supp 2A) Number of merozoites per schizont in PP7-HA:loxP parasites treated with DMSO or rapamycin. Results are the average number of merozoites per schizont of two biological repeats (a minimum of 15 schizonts were counted per replicate). Error bars, SD. Mann–Whitney U test was performed for statistical analysis (ns: not significant).

(Supp 2B) DNA content of DMSO and rapamycin-treated PP7-HA:loxP schizonts. The histograms are representative of two independent experiment. At least 100,000 events were counted per sample. No obvious difference in the DNA content was detected.

(Supp 2C) Combined DIC and fluorescence images from time-lapse video microscopy of DMSO- and RAP-treated PP7-HA:loxP schizonts taken at 5 min (T = 5) and 30 min (T = 30) after release from compound 2. DMSO parasites were pre-treated with Hoechst to stain the nuclei so that they could be viewed simultaneously with the RAP-treated parasites in the same imaging chamber. Schizonts visible in the first frame that rupture over the course of the video are circled in blue (DMSO) or white (RAPA). Scale bar, 10 μm.

(Supp 2D) Invasion rate of DMSO and rapamycin-treated PP7-HA:loxP parasites. The parasitaemia of schizonts incubated with fresh erythrocytes was determined immediately (0 h) and after 4 hours post re-invasion. The invasion rate was determined as the ratio of the parasitaemia at the two time points (4 h/0 h). Three biological replicates were performed in triplicate. Error bars, error of mean. Mann–Whitney U test was performed for statistical analysis (ns: not significant).

**Supplemental Figure 3:**
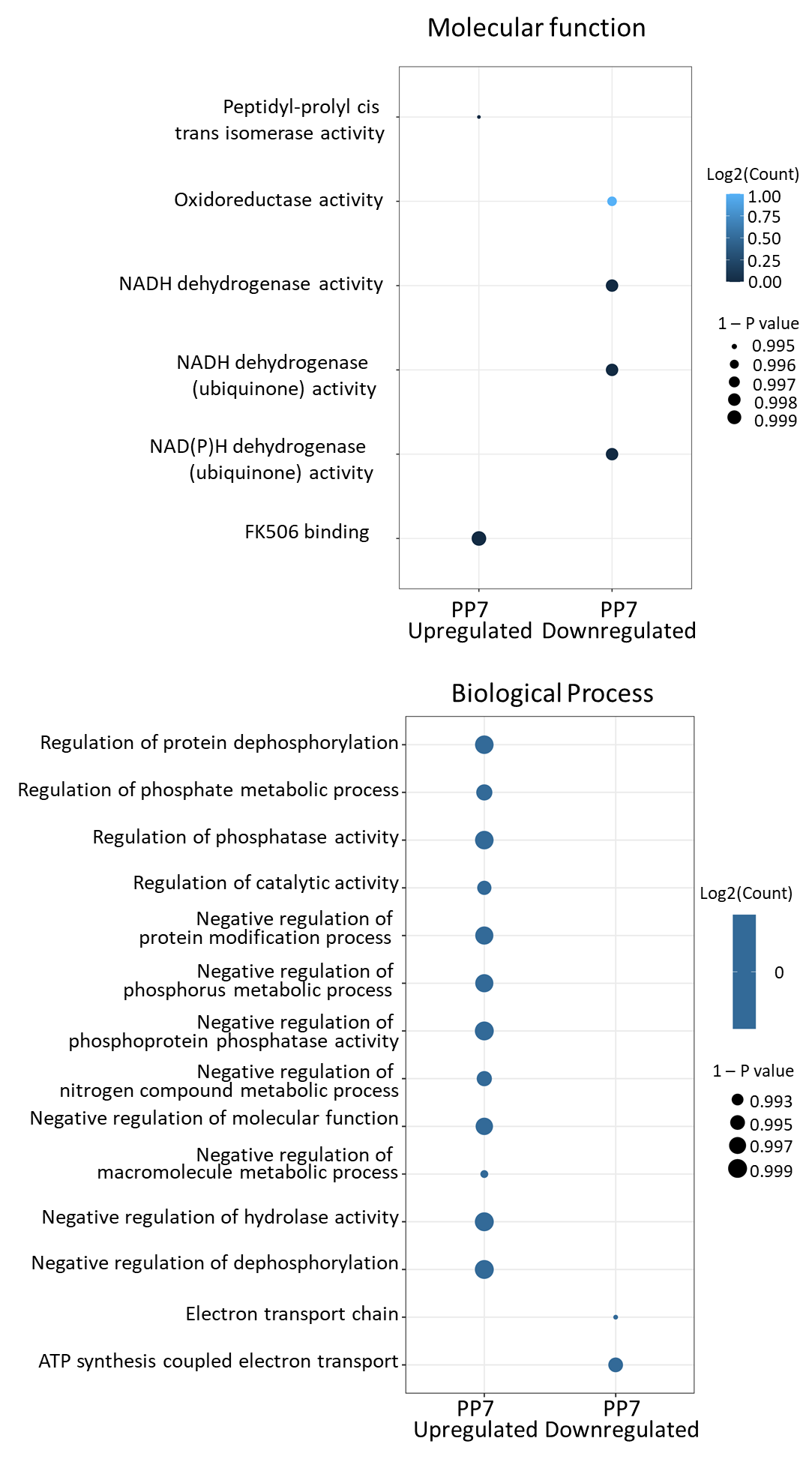

Gene ontology analysis performed on the DMSO and rapamycin-treated PP7-HA:loxP schizonts phosphoproteomes. Gene ontology (GO) enrichment analysis of significantly hyper or hypo phosphorylated proteins in DMSO and rapamycin-treated PP7-HA:loxP schizonts. GO terms were sourced from the PlasmoDB database. The level of significance (1-p value) of the enriched GO term is indicated by the size of the bubble. The colour density indicates the number of differentially expressed proteins (log2 protein count) associated with the GO term.

**Supplementary Figure S4**

(A)

ATATTAGGAGACGTTCATGGACAGTTACATGATGTCTTGTGGCTGTTTAACAGGTTCGGAATTCCGTCCTCAACCAACATCTACATCTTCAACGGTGACATCGCGGATAGGGGTGAAAACGCGACAGAGATCTTTATCCTATTGTTCATCTTTAAACTTAGCTGTAACGACTCTGTGATAATAAACAGAGGCAATCATGAGTGTAGTTACATGAACGAGGTTTACGGCTTCTACAATGAGGTTCTTTCTAAGTATGATAATACGATCTTTGACCTATTTCAAAATATTTTTGAACTACTGGGATTAGCAGTGAATGTACAGAACCAAATATTCGTGGTTCATGGTGGCCTTTCAAGATACCAGGACATAACCCTGAAGGAAATCGACGAGTTAGATAGAAAAAAACAGGAAATTCTTCACCCGGAGCAGTATGAAGACATAGTTATCTTCGATCTGTTGTGGTCTGACCCCCAGAAGAAAGAAGGCATCGGCGGTAATGCGCGTGGGAATAATTGTATAACCTTTGGGCCAGATGTCACAGAAATGTTCTTGAAAAATAATAATCTGGACATTCTTATTAGGTCACATCAGGTGCCAAAGACGCTAAAAGGCATAGAATCCCACCACGAAGGAAAATGTATAACGCTGTTCAGCGCCAGTAATTATTGTAACAAGATAAAAAACTTAGGGGCTGCTATAATATTTAATCAAGATCTTACTTTCGAGGTACAGGAATATATGTCCCCCTCCTTAGAAGTCATTAGGGAAACATTTGAGGAGAACCAGAAACTTAGAGAAAAGGTGTTGCACTGCTCCAAAATAGTGGAACTTGAAAAGAACGAACAGAAAAACAACAACAAGTTGAGCACTGAGGGTCTTATGAATGATATAATCAACTGTTTGTCAACAATCATCTGCAACGAGAAAAATAGTTTATGGAACAATTTGTATAAACAAGATAAAGACAAGAAAGGTGTAGTACATATAAATATCTGGAAGGAAGAGTTGGGTAAACTGAGTAAAGCAAAGAAGGTCCCTTGGATTTACTTGTGTAGAAAGTTAAAAATGATAGAAGACTATCATGTAAATTACAATAATATACTAAGCAGATTCAAAATAAACTATGCGCCAAATGAGAAGTTTCTAAACACAGAATGGAAAAATGAATGTTTCGAGCATCTATATGAGGCTCTTCTAAAAGCCGACTTATCCTTGAGAGAGACTTTAATGGTTTTTGACAAGAATCTGGACGGCAAGGTAAGCTTTGCAGAATTTGAACAGGTTTTAAGGGACCTAAACATAGATTTATCAAATGAACAAATCCGTATTCTTGTCAGGCTAATAAATTCCAACTCCTTATGCAATAATACAAACTTGCAGGAAAACGACAAGATCGACGTAGCGGAGTTTATAGGCAAAATGAGGGTTTGTTACAGACTTAGTATTAACAAGGACTATGTTAACAATGAGAAGATTCAAAAACTGATAGAAACAATCGGCAAACATATATTAAGCGATTCAGCTGATACTGCTAACTACCATTATAAATTCTACGAGGAGAACAACGAGAGACATAATTCCGAGAGAAGAAAGCGTAGCTCCGTAATTAAGTCCGTGGCCCTTTTTCAGAAATTCAAAAACTACGATAACTTCGGTAACGGTTATTTAGACTATAATGACTTCGTCAAGGCGATAAAAAACTTTGACATGAATAAAATTTCAAAAGAAGTCGAGTTCGAGGTAGACGACGATATCCTTATGGAGTTAGCAAAGTCCATAGATATCACAAAATCCAGTAAAATAAATTTTTTGGAGTTTTTGCAAGCTTTCTACGTGGTAAATAAGTCTAAGTACTCATACGTGGACGAGATTTGGTGCCACATCTGCACCGTGATCTATGAAAATAAGGTCGCCCTTAAAAAGTGTATGAAGTACCTTGAGGACACAATGCAGGGGAAAATTACAAGTATTCATTTTCGTTACATTCTTCTAGAGTTGAACAAAATCCTACAGGAGCACAACTTCGAGATGAATAAGCCGCTGACTGACGAGCAAATTGATTTGCTGGCGTATACAGTTGAGACGGACGATAAAGTGGATTATGTAGAGTTCTTCAACAGCTTCAAACCTACATATATATACAGCAATAAT

A ~800-bp 5′ homology region comprising the native gene sequence upstream of the re-codonised region was cloned upstream of the SERA2loxPint. Following transfection of purified schizonts using an AMAXA nucleofector 4D (Lonza) and P3 reagent (Lonza), modified parasites were selected as described previously (Birnbaum et al., 2017).

(B)

TTTTCTTCCCACATTTCGAATAAAACGCGTATGGTGTCCAAGGGCGAGGAAGACAACATGGCATCTTTGCCTGCCACTCACGAGTTGCATATTTTTGGATCAATTAACGGAGTGGATTTCGATATGGTTGGACAAGGGACGGGCAATCCAAATGATGGCTATGAAGAATTAAACTTGAAAAGCACGAAAGGTGATCTTCAGTTCTCCCCATGGATACTTGTCCCACATATTGGCTATGGGTTTCATCAATACTTGCCGTACCCGGATGGGATGTCCCCGTTTCAGGCAGCTATGGTCGACGGTAGCGGTTATCAGGTGCACAGAACAATGCAATTCGAGGATGGGGCCTCCTTGACGGTCAATTACAGGTACACGTATGAGGGAAGTCACATTAAAGGAGAGGCCCAAGTTAAAGGAACAGGTTTTCCTGCCGATGGACCGGTAATGACGAACAGCCTGACTGCCGCAGATTGGTGTAGGTCTAAAAAAACATACCCTAATGACAAGACAATAATTAGCACCTTTAAGTGGAGCTATACAACTGGTAATGGGAAGAGATATAGGTCAACTGCTAGGACTACCTATACCTTTGCGAAGCCAATGGCTGCTAACTATCTTAAAAATCAGCCCATGTACGTTTTCAGGAAAACAGAGCTAAAGCATAGCAAAACAGAATTGAATTTTAAGGAATGGCAAAAGGCTTTTACGGATGTTATGGGAATGGACGAGCTATACAAAGGATCCGGTGGTGGAGAGGGAAGGGGGTCCCTTCTTACATGTGGCGATGTAGAAGAAAATCCGGGGCCTATGGCTCCATTGTCACAGGAAGAAAGCACATTGATAGAGAGAGCCACTGCTACGATAAATAGCATTCCTATCTCAGAGGACTATTCTGTCGCTTCCGCCGCGTTATCTTCAGATGGTCGTATATTTACGGGAGTCAACGTATACCATTTTACTGGTGGACCCTGCGCAGAGTTAGTGGTCCTTGGTACTGCCGCAGCAGCGGCGGCGGGCAACCTTACCTGCATCGTAGCAATCGGAAATGAAAATAGAGGCATATTGTCACCGTGTGGGAGATGCCGTCAGGTTTTATTGGATTTGCACCCCGGCATTAAAGCCATAGTCAAGGACTCTGATGGGCAGCCTACAGCCGTGGGAATCAGAGAATTGCTTCCGTCAGGGTACGTGTGGGAGGGCTAATAAGGTACCACCTAACAACTTTATGTCGACTC

Donor sequences were constructed by cloning a synthetic DNA fragment encoding mNeon green and the downstream T2A and BSD sequences (IDT) into pDCIn (DiCre induction) plasmid that had been linearised by digestion with MluI and KpnI (Patel et al., 2019).

**Supplemental Movie 1:**

Both DMSO (Hoechst) and RAPA (DIC) treated parasites egress normally. Percentage of egress: DMSO: 36.5% (23/63), RAPA: 35.8% (33/92). Scale bars, 10 μm.

### Supplemental Table 1:

### Proteins identified from pull down experiments of PP7-HA:loxP and 3D7DiCre in P. falciparum schizonts.

### Supplemental Table 2:

### Phosphoproteome analyses of PP7-HA:loxP and 3D7DiCre in P. falciparum schizonts.
